# A chromatin accessibility map of pea aphid brain and embryo identifies tissue-specific regulatory elements

**DOI:** 10.64898/2026.05.14.725175

**Authors:** Xiaomi Liu, Jennifer A. Brisson

## Abstract

The pea aphid (*Acyrthosiphon pisum*) is an important model organism for studying complex biological traits, including wing polyphenism and host-symbiont interactions, yet its regulatory genomic landscape remains largely uncharacterized. Here we present the first genome-wide chromatin accessibility map of the pea aphid, generated using the assay for transposase-accessible chromatin followed by sequencing (ATAC-seq). We profiled open chromatin regions (OCRs) in adult brains and late-stage embryos from winged and wingless morphs maintained under solitary or crowded conditions. We also paired ATAC-seq with RNA-seq in embryonic samples to examine the relationship between chromatin accessibility and gene expression. Libraries showed a high abundance of reads from the aphid endosymbionts *Spiroplasma* and *Buchnera*, reflecting preferential Tn5 transposase insertion into nucleosome-free bacterial DNA. After computational removal of these reads, the remaining aphid-mapping libraries displayed hallmarks of high-quality ATAC-seq data. We identified a consensus set of 37,127 OCRs enriched at promoters and distal regulatory elements, with substantial overlap with computationally predicted enhancers and enrichment for transcription factor binding motifs. Tissue identity was the dominant driver of chromatin variation, accounting for 85% of variance along the first principal component, with 19,513 differentially accessible regions distinguishing brain from embryo samples. By contrast, differences associated with wing morph or crowding treatment were modest. Promoter accessibility was significantly and positively correlated with gene expression genome-wide. Together, these data constitute a foundational regulatory genomics resource for the pea aphid and establish a framework for mechanistic studies of gene regulation in this ecologically and economically important insect.

## Introduction

Gene regulation is shaped by chromatin organization, and in particular by the physical accessibility of genomic DNA to transcription factors and the transcriptional machinery. Chromatin is open at active regulatory elements such as promoters, enhancers, and other cis-regulatory modules (Klemm et al. 2019), and open chromatin is strongly predictive of transcriptional activity (Thurman et al. 2012). Mapping open chromatin regions (OCRs) across tissues and developmental stages, therefore, provides a window into a genome’s regulatory architecture. The assay for transposase-accessible chromatin followed by sequencing (ATAC-seq) has become the method of choice for this purpose (Buenrostro et al. 2013). By exploiting the preferential insertion of the Tn5 transposase into more open, nucleosome-depleted DNA, ATAC-seq enables genome-wide accessibility profiling from small numbers of cells or nuclei, making it particularly well-suited to organisms and tissues where sample material is scarce.

ATAC-seq has been applied across a variety of insects, including *Drosophila melanogaster* (Blythe and Wieschaus 2016), *Apis mellifera* (Lowe et al. 2022), and a growing number of other species [*e.g*., the mayfly (Pallarès-Albanell et al. 2024)], revealing how tissue identity, developmental stage, and environmental experience shape the chromatin landscape. These studies consistently find that many OCRs are tissue-specific, that promoter accessibility correlates broadly with gene expression, and that distal regulatory elements contribute to transcriptional differences between cell types and conditions. Despite this progress, chromatin accessibility remains uncharacterized in many insects with otherwise strong biological and genomic resources (Erdoğan et al. 2025).

The pea aphid (*Acyrthosiphon pisum*) is one such species. Pea aphids have long served as a model for studying the genetic and developmental bases of complex traits, particularly wing plasticity (Brisson et al. 2026) and host-symbiont interactions (Smith and Moran 2020). This work has been aided by the early sequencing of the pea aphid genome (Consortium 2010), along with refined genome assemblies (Y. Li et al. 2019; Mathers et al. 2021) and annotations (Deem and Brisson 2026). The only existing open chromatin data in this species come from a FAIRE-seq study of whole male and female individuals, which identified sex-specific differences in X chromosome accessibility and linked them to a dosage compensation mechanism (Richard et al. 2017). While that study established that chromatin accessibility is biologically meaningful in pea aphids, whole-body FAIRE-seq data are not well-suited to resolving tissue-specific regulatory programs or producing a genome-wide annotation of cis-regulatory elements. A tissue-resolved chromatin accessibility map, which would be the foundation for mechanistic studies of gene regulation in this species, has not been generated.

Here we present the first genome-wide tissue-specific chromatin accessibility map of the pea aphid. We used ATAC-Seq to profile the OCRs of two tissues central to aphid development and behavior: the adult brain and the late-stage embryo. We further paired the ATAC-seq with RNA-seq of the embryonic tissues, using the combined data to examine the relationship between chromatin accessibility and gene expression. Tissue identity emerged as the dominant driver of chromatin variation, with brain and embryo exhibiting extensively divergent accessibility profiles that reflect their distinct regulatory programs. These data provide a foundational regulatory genomics resource for the pea aphid and a framework for future studies of gene regulation in aphids.

## Methods

### Pea aphid stock rearing

First instars of line SSC3 were transferred to *Vicia faba* seedlings (Improved Long Pod, Harris Seeds) every 11-13 days at a transfer density of four per plant. Each plant was planted in a single plastic plant pot (S17648, Fisher Scientific), enclosed in a cylindrical cage cut from clear PETG tubes (VISIPAK), and sealed with fine-mesh netting (Noseeum netting, 117 inches, Barre Army Navy Store). Stocks were maintained at a humidity of 30 ± 10%, temperature of 19 ± 3.5°C, 16h light to 8h dark cycle. Pea aphid stocks reproduced asexually and viviparously under these rearing conditions. Stocks were reared under these conditions for multiple generations to eliminate transgenerational density effects before experiments.

### Sample generation (winged or wingless, crowded or solitary)

We used two treatments - crowded or solitary - and two morphs - winged or wingless - for our collections. To generate wingless aphids, day 12, wingless adult aphids were placed individually on plant seedlings for 24h, after which the adults were removed, and the offspring were kept on the plants to develop into adults. To generate winged aphids, day 12, wingless adult aphids were collected and placed in 35 mm petri dishes at a crowded density of 12 per dish to induce winged offspring. Then, crowded aphids were placed individually on plant seedlings for 24h, after which the adults were removed, and the offspring were kept on the plants to develop into adults.

At adulthood, we collected groups of winged or wingless aphids that we used for sample collection. We subjected each morph to either a crowding (12 individuals in a 35 mm Petri dish) or a solitary condition (individually placed in 35 mm Petri dishes) for 24 h. Treated aphids were then either transferred onto plants for reproduction for 24h to validate the treatment effect on wing induction or dissected for sequencing.

### Tissue collection

We washed aphids two times in 500 ul cold 0.1% PBT to remove bacteria and dirt, and then in 500 ul 1X cold PBS twice to remove any PBT residue. We transferred the washed aphids to a glass well dish filled with 500 ul cold 1X PBS and carefully dissected out the brain and fused ventral nerve cord (hereafter referred to as brain for simplicity) and embryos as needed. For the brain samples, we removed the salivary glands that are usually attached.

Embryos at stages 18-19, with a length of around 1000 μm, were used (Miura et al. 2003). We washed dissected tissue in clean 1X cold PBS two more times and then stored it in 1X PBS on ice until all dissections were completed. Each biological replicate was generated from three individual adult aphids. For each replicate, one brain and one embryo were dissected from each adult (three brains or three embryos total) and pooled.

Because embryos derived from wingless adults predominantly develop into winged morphs, whereas embryos derived from winged adults predominantly develop into wingless morphs (Figure S1), the terms “wingless-derived” and “winged-destined” embryos, as well as “winged-derived” and “wingless-destined” embryos, are used interchangeably throughout this study.

### Reagent preparation

All reagents were prepared fresh and maintained ice-cold throughout the procedure. The resuspension buffer was prepared by combining 100 µL Tris-HCl (pH 7.5), 30 µL 1 M MgCl₂, and 20 µL 5 M NaCl with nuclease-free water to a final volume of 10 mL. The WASH buffer was prepared by mixing 990 µL RSB with 10 µL 10% Tween-20. The digitonin lysis buffer was prepared by combining 970 µL RSB with 10 µL 10% Tween-20, 10 µL NP-40, and 10 µL 1% digitonin.

### Nuclei isolation from brain tissue

Dissected brains were processed immediately for nuclei isolation. Briefly, 10 µL of 0.1% digitonin lysis buffer was added to a 1.7 mL microcentrifuge tube. Dissected brains were transferred onto the tip of a sterile plastic pestle and gently homogenized in the tube by pressing the pestle against the tube wall. Homogenization was performed slowly to minimize bubble formation. The pestle was rinsed with an additional 90 µL of 0.1% digitonin lysis buffer, and the wash was collected into the same tube, yielding a total volume of 100 µL. Care was taken to ensure no visible tissue remained on the pestle before disposal. The lysate was gently mixed by pipetting, avoiding air bubble introduction, and incubated on ice for 2 min. Following lysis, 1 mL of wash buffer was immediately added, and the sample was mixed gently by pipetting. The sample was centrifuged at 200 × g for 5 min at 4 °C. After centrifugation, 900 µL of the supernatant was carefully removed using a 1000 µL pipette. The sample was centrifuged again at 200 × g for 3 min at 4 °C to stabilize the nuclei pellet. The remaining supernatant was then gently removed using a 200 µL pipette, leaving approximately 8-10 µL of liquid to avoid disturbing the nuclei pellet.

### Nuclei isolation from embryo tissue

Dissected embryos were processed following completion of brain sample processing. A total of 300 µL of 0.1% digitonin lysis buffer was added to a pre-chilled glass homogenizer. Embryos were carefully transferred to the prechilled glass homogenizer pestle and gently homogenized by slowly pressing and twisting the pestle three times so that no visible tissue remained. Homogenization was performed carefully to minimize air bubble formation. The pestle was slowly withdrawn and rinsed with the homogenate to ensure complete sample recovery, confirming that no visible tissue remained attached. The homogenate was gently mixed by pipetting within the homogenizer. Samples were incubated on ice for a total of 2 min from the start of lysis. Following incubation, 700 µL of wash buffer was immediately added, and the sample was mixed gently by pipetting. The lysate was then transferred to a 1.7 mL microcentrifuge tube and centrifuged at 200 × g for 5 min at 4 °C. After centrifugation, 900 µL of the supernatant was carefully removed using a 1000 µL pipette. The sample was centrifuged again at 200 × g for 3 min at 4 °C to stabilize the nuclei pellet. The remaining supernatant was then gently removed using a 200 µL pipette, leaving approximately 8-10 µL of liquid to avoid disturbing the nuclei pellet.

### RNA-seq

Nuclei were resuspended in the remaining 10 µL of buffer. A 2 µL aliquot of the nuclei suspension was transferred into 300 µL of Trizol (Invitrogen). Samples were immediately stored at −80°C until further processing. RNA was extracted following the standard Trizol protocol. Library preparation and sequencing were performed at the University of Rochester Genomics Research Center. Libraries were sequenced on an Illumina NovaSeq X Plus platform using 150 bp paired-end reads.

### ATAC-seq

Tagmentation and library preparation were performed following the ATAC-seq protocol as described by Buenrostro et al. 2013 with minor modifications as noted here. Nuclei were resuspended in 50 µL transposition reaction mix containing 25 µL Tagment DNA (TD) buffer, 0.5 µL 10% Tween-20, 0.5 µL 1% digitonin, 1× PBS, Tagment DNA Enzyme (TDE1) (0.41 µL for brain samples; 2.5 µL for embryo samples), and nuclease-free water to volume (7.09 µL for brain samples; 5 µL for embryo samples). TDE1 enzyme volume was adjusted for brain samples to account for the low nuclei input. Reactions were incubated at 37 °C for 30 min with a brief and gentle vortex every 5 min to prevent nuclei settling. Tagmentation was stopped by adding 300 µL DNA binding buffer (Zymo DNA Clean & Concentrator-5 kit). DNA was purified according to the manufacturer’s instructions and eluted in 11 µL nuclease-free water.

Initial PCR amplification was performed in a 50 µL reaction containing 10 µL transposed DNA, 1 µL nuclease-free water, 6.25 µL of 10 µM Nextera PCR Primer 1, 6.25 µL of 10 µM Nextera PCR Primer 2, 1.5 µL EvaGreen dye (Biotium), and 25 µL NEBNext High-Fidelity 2× PCR Master Mix (NEB). PCR cycling conditions were as follows: 72 °C for 5 min; 98 °C for 30 s; followed by 5 cycles of 98 °C for 10 s, 63 °C for 30 s, and 72 °C for 1 min; hold at 4 °C.

To determine the optimal number of additional amplification cycles, a qPCR-based approach was used. A 15 µL reaction containing 5 µL pre-amplified product, 3.45 µL nuclease-free water, 0.625 µL each of 10 µM Nextera PCR Primer 1 and 2, 0.3 µL EvaGreen dye (Biotium), and 5 µL NEBNext High-Fidelity 2× PCR Master Mix (NEB) was subjected to real-time PCR on a BioRad CFX96. Additional cycles were determined based on the cycle number corresponding to one-quarter of the maximum fluorescence intensity. The remaining 45 µL of pre-amplified library was then amplified using the determined number of cycles under the following conditions: 98 °C for 30 s; followed by N cycles of 98 °C for 10 s, 63 °C for 30 s, and 72 °C for 1 min; hold at 4 °C. PCR products were purified by adding 225 µL DNA binding buffer (Zymo DNA Clean & Concentrator-5 kit) and processed according to the manufacturer’s instructions. Purified libraries were submitted to the University of Rochester Genomics Research Center for sequencing. Prior to sequencing, fragment size distribution was assessed with a Bioanalyzer. Samples were sequenced on an Illumina NovaSeq X Plus platform with 150 bp paired-end reads.

### ATAC-seq data quality assessment

Raw sequencing reads were processed using fastp 1.0.1 (Chen et al. 2018) to remove adaptors, poly-G tails, and low-quality reads, with quality control report generation enabled. Cleaned reads were aligned to the pea aphid reference genome v3.0 (Y. Li et al. 2019) using bowtie2 2.3.5.1 in -sensitive-local mode (Langmead and Salzberg 2012). Library composition and potential contamination were assessed using Kraken2 2.1.5 (Wood et al. 2019). According to the Kraken2 result, two additional custom reference genomes were constructed by downloading 6 genomes of *Spiroplasma ixodetis* (Taxid 2141) and 176 genomes of *Buchnera aphidicola* (Taxid 9) from NCBI, which were concatenated into separate composite genomes for targeted mapping. Duplicated reads were marked using Picard 3.4.0 (https://broadinstitute.github.io/picard/) MarkDuplicates function within the GATK framework (McKenna et al. 2010). Aligned reads were further filtered with samtools 1.21 (H. Li et al. 2009) with the following criteria: mapping quality ≥30, properly paired reads, and exclusion of secondary, supplementary, and duplicate alignments (-q 30 -f 2 -F 1804). Reads mapping to the mitochondrial genome were removed prior to downstream analyses. Fragment size distributions were estimated using Picard 3.4.0 (https://broadinstitute.github.io/picard/) CollectInsertSizeMetrics within the GATK framework (McKenna et al. 2010). Tn5 insertion bias was corrected using deepTools 3.5.1 (Ramírez et al. 2016) alignmentSieve with the --ATACshift option. Fragments were further categorized as nucleosome-free fragments (fragment size <= 100) or mono-nucleosome fragments (180 <= fragment size <= 247). Genome-wide coverage tracks were generated for nucleosome free and mono-nucleosome fragments separately using deepTools 3.5.1 (Ramírez et al. 2016) bamCoverage as reads per genomic content (--normalizeUsing RPGC --effectiveGenomeSize 541208503 binSize 10). Chromatin accessibility quality was assessed by calculating enrichment around transcription start sites (TSS ±2 kb) extracted from aphid annotation (Deem and Brisson 2026) using deepTools 3.5.1 (Ramírez et al. 2016) computeMatrix.

### ATAC-seq peak calling and annotation

Reads aligned to the pea aphid genome were used to call open chromatin regions (hereafter referred to as OCRs) in each sample using MACS 2.1.1 (Zhang et al. 2008) with paired-end mode, effective genome size 5.4 x 10^8^, non-model organism, and adjusted p-value cutoff 0.05. OCRs were further filtered to retain high-confidence regions based on q-value <= 0.01 (FDR-controlled significance threshold) and MACS2 narrow peak score >= 2. Filtered OCRs from all 24 samples were combined into a consensus OCR set using DiffBind 3.12.0 (Stark and Brown, 2011) Only OCRs present in at least three samples were retained to ensure reproducibility. Read counts over the consensus OCRs were quantified using featureCounts 2.0.3 (Liao et al. 2014) for downstream statistical analyses. OCRs were classified as 1) intergenic, 2) promoter-like (2 kb upstream and 200 bp downstream of the TSS, hereafter referred to as promoter-like OCRs), 3) exonic, and 4) intronic based on their genomic context using the modified reference pea aphid genome annotation described in (Deem and Brisson 2026).

### RNA-seq analysis

Raw reads were processed with fastp 1.0.1 (Chen et al. 2018) to remove adaptors and low-quality reads, with quality control report generation enabled. Cleaned reads were mapped back to pea aphid genome v3.0 (Y. Li et al. 2019) using hisat2 2.1.0 (Kim et al. 2019) with the following parameters: -D 20 -R 3 --score-min L,0,-0.6 -N 1 -k 10. Read counts over exons were calculated with featureCounts 2.0.3 (Liao et al. 2014) using the pea aphid annotation (Deem and Brisson 2026) and summarized at the gene level for statistical comparisons.

### DAR and DEG analysis

Differential accessibility (ATAC-seq) and differential expression (RNA-seq) analyses were performed using the R package DESeq2 (Love et al. 2014) with default settings; raw count matrices were normalized using DESeq2’s median-of-ratios method, and statistical significance was assessed using the Wald test. Resulting p-values were adjusted for multiple testing using the Benjamini-Hochberg procedure. Principal component analysis (PCA) was performed on DESeq2 variance-stabilized count data (generated with blind = FALSE, which incorporates the experimental design during transformation to preserve condition-associated variation). PCA was conducted using the most variable features. Significance thresholds for differentially accessible regions (DARs) and differentially expressed genes (DEGs) were adjusted p-value (FDR) <= 0.05 and fold change >= 1.5. All comparisons were performed by subsetting the relevant samples, followed by differential analysis using a single-factor design.

Nearest genes around the DARs were retrieved with bedtools v2.31.1 (Quinlan and Hall 2010). One DAR might be located in the middle of a gene, or multiple genes if there are multiple annotations in the same place. Gene homology annotations were obtained from the reference genome annotation file (Deem and Brisson 2026) and used to map genes to gene ontology terms based on NCBI annotations. We performed gene-level gene ontology enrichment analysis using all nearest genes around OCRs, and gene set enrichment analysis (min gene set size = 10, max gene set size = 300) with the R package clusterProfiler (Yu et al. 2012; Wu et al. 2021).

### Motif enrichment

Motif enrichment analysis was performed using HOMER findMotifsGenome.pl (http://homer.ucsd.edu/homer/ngs/peakMotifs.html). Both consensus OCRs and tissue-specific DARs were analyzed. For consensus OCRs, background regions were generated by randomly sampling genomic regions with -size given option using HOMER’s internal background selection. For tissue-specific DARs, non-differentially accessible OCRs were used as the background set to control for baseline chromatin accessibility.

### Enhancer prediction

To investigate the regulatory architecture of the pea aphid genome, putative enhancers were predicted using SCRMshaw_HD (Kazemian et al. 2011; Asma and Halfon 2019). All 48 precompiled training datasets (https://github.com/HalfonLab/dmel_training_sets) were used, and three prediction algorithms (hexMCD, IMM, and PAC) were implemented in SCRMshaw_HD to maximize sensitivity.

Enhancer predictions (500 bp in length) generated from all training datasets and algorithms were combined using a custom script. Within each prediction method, candidates were ranked by score, and the top 20% were retained to enrich for high-confidence predictions. Retained predictions from all methods were then merged across methods, and training sets, and overlapping regions were consolidated to generate a non-redundant set of consensus predicted enhancers.

### ATAC-seq and RNA-seq paired analysis

Genes were classified based on the presence or absence of promoter-like OCRs. Gene expression levels between these two groups were compared using the Wilcoxon rank-sum test. We used a pairwise Spearman’s rank correlation test to compare chromatin accessibility in the promoter region and gene expression level (n = 6). To evaluate global concordance between chromatin accessibility and gene expression, correlation coefficients were standardized (Z-score transformation) and compared to a null distribution generated by permutation testing (500 randomizations), in which sample labels were shuffled.

## Results

### ATAC-seq libraries show enrichment of nucleosome-free DNA and strong signal at transcription start sites

To evaluate ATAC-seq data quality, we examined standard mapping metrics for all 24 samples. Initial alignment to the pea aphid genome yielded a low mapping rate (13% to 29%), suggesting substantial contamination from non-aphid DNA (Table S1). Taxonomic classification with Kraken2 (Wood et al. 2019) showed that a large fraction of reads originated from the aphid endosymbionts *Spiroplasma ixodetis* and *Buchnera aphidicola*. Consistent with this, reads from brain samples (n = 12) predominantly mapped to *Spiroplasma ixodetis* (41-77%), whereas reads from embryo samples (n = 12) predominantly mapped to *Buchnera aphidicola* (65-80%; Table S1). When these endosymbiont genomes were included, overall mapping rates increased to 69-97%. These results indicate that the low initial mapping rates primarily reflect preferential enrichment of symbiont-derived reads, likely due to higher chromatin accessibility in bacteria (*e.g*., lack of nucleosomes), which facilitates Tn5 insertion, rather than poor DNA quality (*e.g*., degradation).

After correcting for symbiont contamination, we examined fragment size distributions for reads mapped to the aphid genome. The libraries are dominated by nucleosome-free fragments, while mono- and di-nucleosome fragments (∼200 bp and ∼400 bp) are weakly represented (Figure 1A). Consistent with this, nucleosome-free fragments show strong enrichment at transcription start sites (TSSs; Figure 1B), indicating effective capture of accessible chromatin. Mono-nucleosome fragments show enrichment around TSSs with only a shallow depletion at the TSS, indicating that while accessibility signals are captured, canonical nucleosome positioning patterns are weakly resolved. Together, these results indicate that the libraries robustly capture open chromatin but are not well suited for analyses requiring high-resolution nucleosome phasing (*e.g*., nucleosome positioning); accordingly, we did not pursue such analyses.

**Figure 1.**
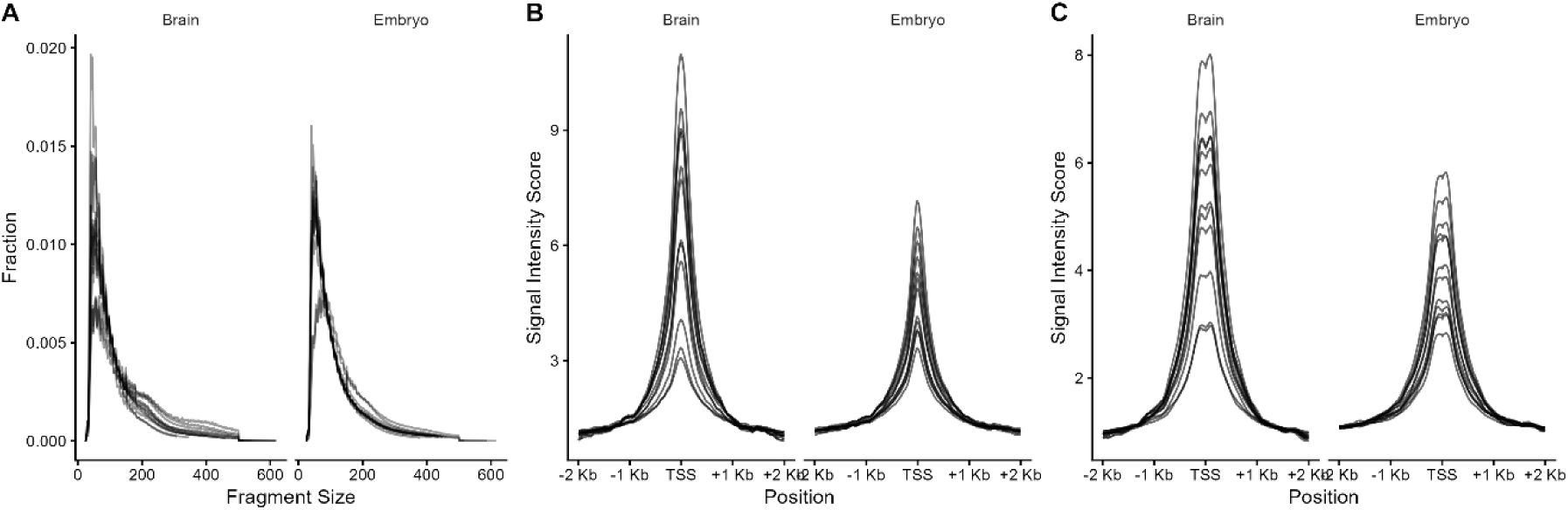
ATAC-seq quality control. (A) Fragment size distribution of ATAC-seq libraries. (B) Signal intensity scores around TSS for nucleosome-free fragments (fragment size <=100 bp). (C) Signal intensity scores around TSS for mono-nucleosome fragments (180 <= fragment size <= 247 bp). For A-C, each line corresponds to one library. Brain samples are shown on the left within each panel, while embryo samples are shown on the right.

### OCR characterization reveals widespread enrichment of accessible chromatin at regulatory elements

To compare chromatin accessibility across samples, we identified OCRs for each library and constructed a consensus OCR annotation set. After score and adjusted p-value filtering, individual libraries yielded 9,351-10,138 OCRs, and the final consensus set contained 37,127 OCRs. OCR lengths ranged from 108 bp to 3,841 bp, with a mean of 589 bp (Figure 2A).

**Figure 2.**
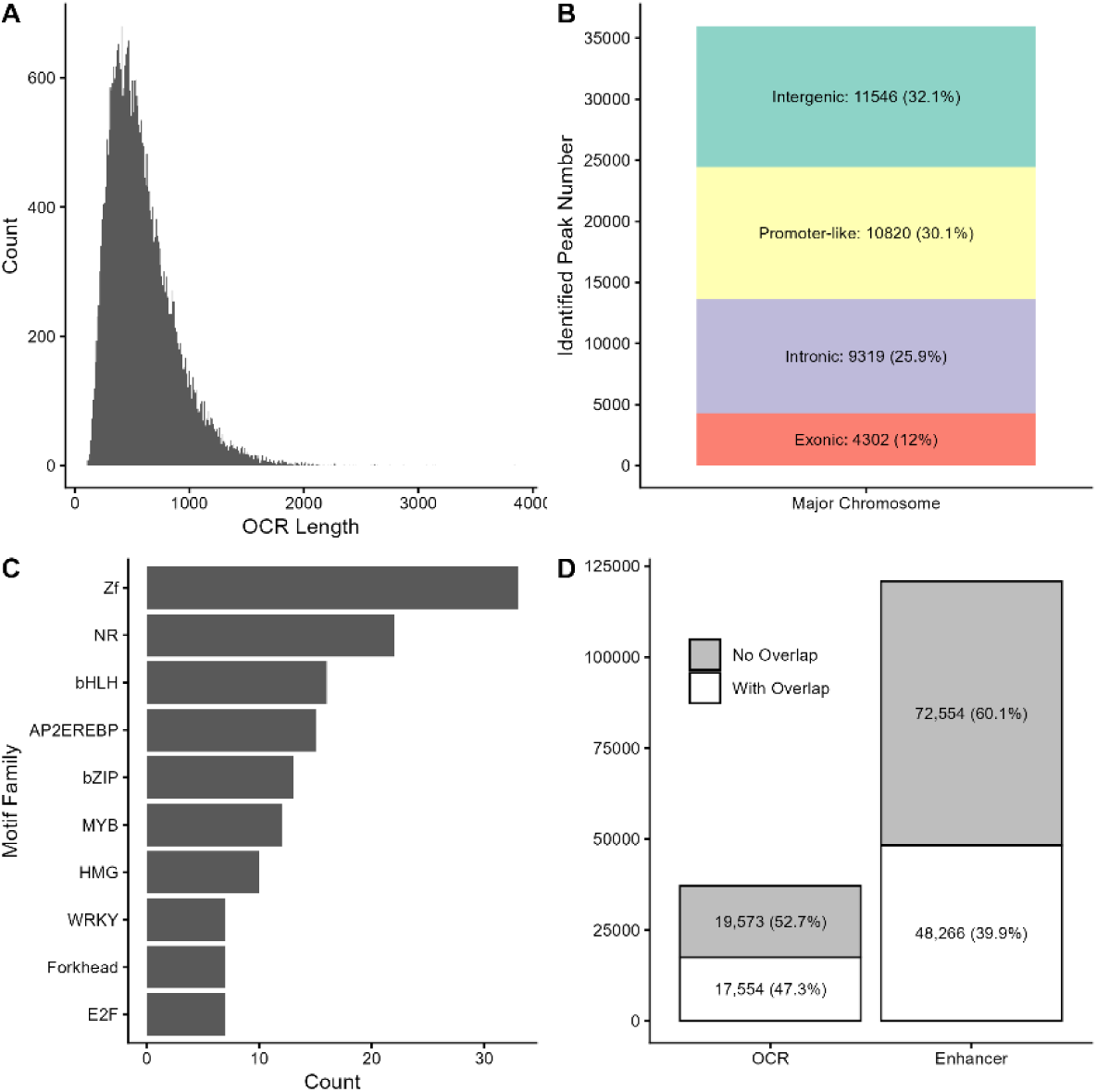
Open chromatin region (OCR) characteristics. (A) Histogram with 50 bp bin width showing the distribution of OCR lengths for the consensus peak set. (B) Genomic annotation of OCRs, categorized as intergenic, promoter (−2 kb to +200 bp relative to TSS), intronic, or exonic. (C) Top 10 enriched transcription factor motif families identified in OCRs, using randomly selected genomic regions as background. (D) Overlap between OCRs and computationally predicted enhancers using a ≥20% OCR-length overlap cutoff.

We characterized the genomic distribution of these OCRs to assess their regulatory context. We annotated OCRs into four categories: intergenic, promoter (using the commonly used demarcation of −2 kb to +200 bp relative to TSS), intronic, and exonic (Figure 2B). Intergenic regions comprised the largest fraction, consistent with their genome-wide abundance. Promoter-associated OCRs were the second-largest category, aligning with the strong TSS enrichment observed earlier and indicating effective capture of regulatory regions.

To further evaluate regulatory potential, we performed motif enrichment analysis of these OCRs. We identified 201 overrepresented transcription factor binding motifs, such as zinc finger (ZF), nuclear receptor (NR), and basic helix-loop-helix (bHLH) families (Figure 2C, Table S2), supporting the trans-regulatory relevance of the detected OCRs.

To assess the distal regulation potential of these OCRs, we examined overlaps between OCRs and computationally predicted enhancers (≥20% OCR length overlap used as a cutoff). We identified 120,820 predicted enhancers (median length = 500 bp, mean length = 711.5 bp. A total of 17,554 OCRs (47.3% of all OCRs) overlapped at least one predicted enhancer (Figure 2D). Given their sometimes large size, OCRs can potentially overlap multiple enhancers.

Reciprocally, 48,266 predicted enhancers (39.9% of all enhancers) overlapped OCRs. These reciprocal overlaps support the regulatory relevance of the identified OCRs and the predicted enhancer set.

### ATAC-seq differential accessibility analysis reveals chromatin variation across tissue, phenotype, and treatment

Having established the overall quality and regulatory relevance of the ATAC-seq data, we examined differential chromatin accessibility across tissues, wing phenotypes, and treatments. Our dataset consisted of brain and embryo samples derived from winged and wingless aphids that had been treated with either 24h solitary or crowded conditions (Figure 3A).

**Figure 3.**
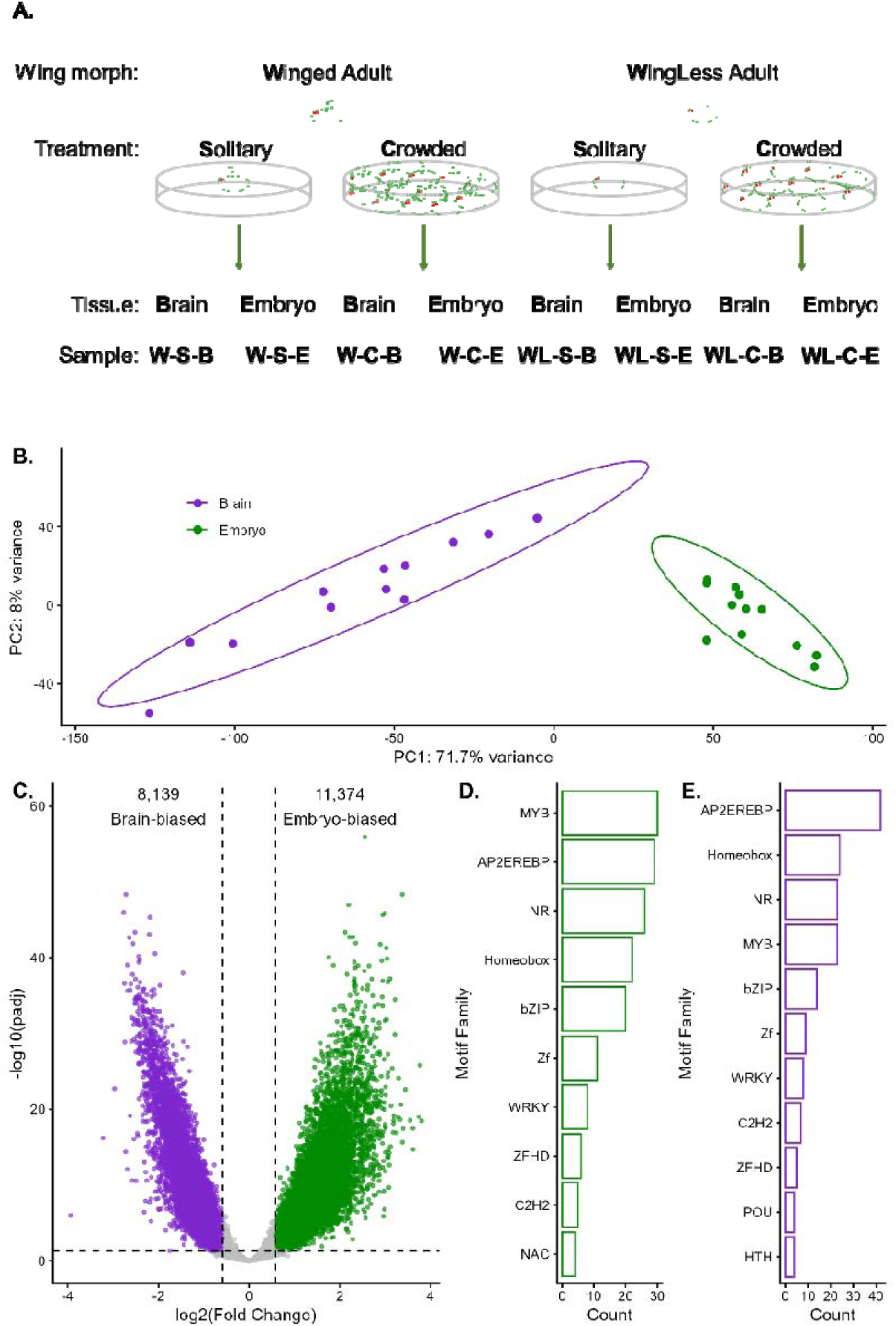
Tissue-specific chromatin accessibility profiles. (A) Sample collection design. (B) Principal component analysis (PCA) of 24 ATAC-seq samples, with the top 10,000 variable OCRs used for analysis. Points are colored by tissue: brain (purple) and embryo (green). (C) Volcano plot of differentially accessible regions (DARs) between tissues. Embryo-biased DARs are shown in green, and brain-biased DARs in purple. (D-E) Top 10 enriched transcription factor motif families identified from (D) embryo-biased DARs, and (E) brain-biased DARs. Non-differentially accessible regions were used as background for enrichment analysis.

We assessed chromatin accessibility differences across treatment (crowded vs. solitary), wing phenotype (winged vs. wingless), and tissue (brain vs. embryo) using pairwise comparisons. Tissue had the dominant effect. Principal component analysis (PCA) showed clear separation by tissue along PC1 (Figure 3B), explaining 71.7% of the total variance, with tissue alone accounting for 85.0% (Table S3). Consistent with this, tissue contrasts (12 vs. 12) identified extensive differential accessibility: of 37,127 OCRs analyzed, 11,374 were more accessible in embryos and 8,139 in brains (Figure 3C). These large differences likely reflect fundamental regulatory divergence between these distinct tissue types.

To assess the regulatory relevance of these tissue-associated differentially accessible regions (DARs), we performed motif enrichment analysis using HOMER (http://homer.ucsd.edu/homer/ngs/peakMotifs.html). We identified 206 enriched motifs in embryo-biased DARs, and 193 in brain-biased DARs. Developmentally relevant transcription factor families were prominently enriched. Homeobox motifs ranked among the top hits across comparisons (Figure 3D-E, Table S4, S5), and zinc finger homeodomain (ZFHD) motifs were also consistently enriched. Their high ranking relative to background (not relative to significantly differentially accessible OCRs) supports that these DARs capture biologically meaningful regulatory differences associated with tissue identity and development.

To assess the potential biological functions of genes associated with these tissue-specific DARs, we retrieved the nearest genes and performed gene-level gene ontology enrichment analysis. The majority of DARs have genes within 10 kb (Figure S2). For genes around embryo-biased DARs, gene ontology enrichment terms are primarily associated with metabolism (Figure 4A), while genes around brain-biased DARs are associated with neurons and neuronal activity (Figure 4B). See Figure 4A-B and Table S6, S7.

**Figure 4.**
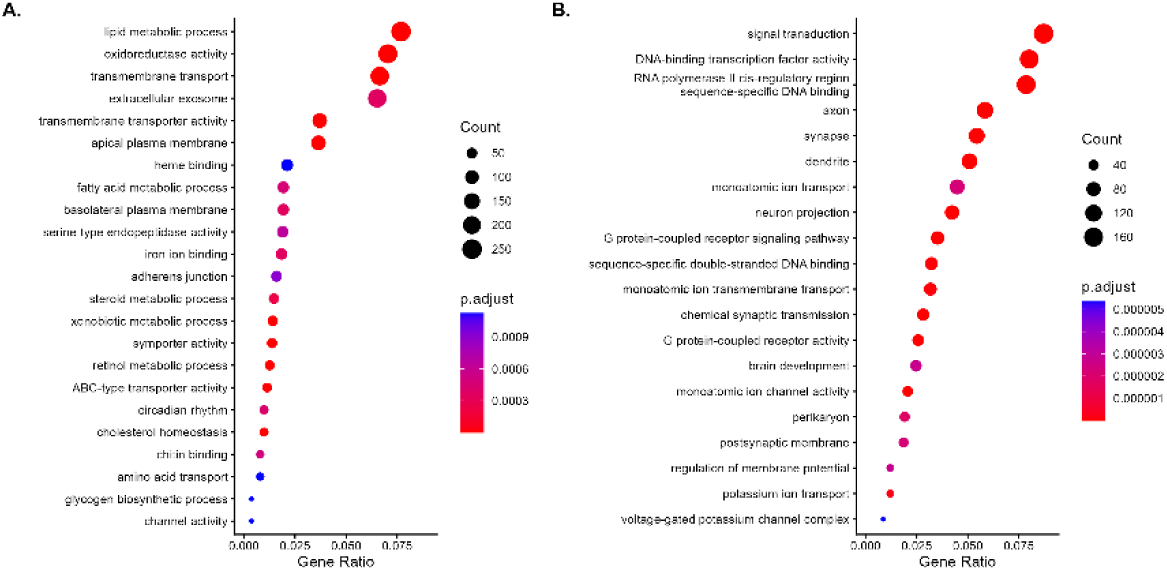
Gene ontology (GO) term for genes around tissue-specific DARs. (A) Top enriched GO term for genes within 10kb of embryo-biased DARs. (B) Top enriched GO term for genes within 10kb of brain-biased DARs.

We performed pairwise comparisons contrasting wing morphs and treatment, nested within each tissue type, to account for the confounding effect of tissue type. In contrast to the substantial differences between tissues, chromatin differences associated with wing phenotype were modest. Comparing the six winged-destined embryo samples to the six wingless-destined embryo samples identified 975 DARs (Figure S1A). However, within-treatment, between morph comparisons (three versus three for each) showed no significant DARs for solitary conditions (Figure S1B) and only 11 for crowded conditions (Figure S1C). The larger scale of DARs in the treatment-comparison combined analysis likely reflects, at least in part, increased statistical power (six versus six compared to three versus three).

Brain samples showed even fewer differences. Comparing the six brains from adult wingless individuals compared to six brains from winged females resulted in only four DARs (Figure S1G). We observed no significant differences within treatment groups (Figure S1H, I) or across treatment conditions (Figure S1 D-F, J-L). This is consistent with the phenotype validation data, where treatment shows no detectable treatment effect for offspring phenotype in winged morphs and only a mild effect in wingless morphs (Figure S1).

### RNA-seq analysis of crowded embryo samples reveals modest but functionally biased transcriptional differences associated with wing morph fate

To investigate transcriptomic differences associated with eventual wing morph fate, we focused on embryo samples collected from winged or wingless mothers following a crowded treatment. This choice was driven by the ATAC-seq results: treatment contrasts did not yield detectable differential accessibility, whereas wing-phenotype contrasts produced modest but detectable signals, and crowded embryos showed more visible differences compared to solitary embryos (Figure S1, A-C). We therefore sequenced RNA from three embryo samples derived from crowded winged adults (which contain wingless-destined embryos) and three from crowded wingless adults (which contain winged-destined embryos). These RNA-seq samples were paired with ATAC-seq data, as both were generated from the same pooled nuclei samples.

Differential expression analysis revealed modest transcriptomic divergence between the two groups, mirroring the mild differences in chromatin accessibility observed in the ATAC-seq analysis. PCA showed only subtle separation by wing fate (Figure 4A), with wing morph contributing the most to PC1 and PC3 (34.4% to PC1, which explains 46.5% of total variance and 32.6% to PC3, which explains 14.3% of total variance, Table S8). This subtle clustering is consistent with a small number of differentially expressed genes (DEGs n = 62; Figure 4B). Of these, 39 genes had higher expression in crowded winged-derived (wingless-destined) embryo samples (n = 3), and 23 showed higher expression in crowded wingless-derived (winged-destined) embryo samples (n = 3).

To assess the functional implications of transcriptional differences between different morph-destined embryos, we performed gene set enrichment analysis, which identified 162 significantly enriched pathways (Figure 5D, Table S9). Most pathways were wingless-derived/winged-destined embryo-biased, suggesting asymmetric activation of transcriptional programs. Pathways enriched in winged-derived embryos are mostly related to sensory and olfactory functions, while pathways enriched in wingless-derived embryos are mostly related to lipid metabolism, transmembrane transporter activity, DNA replication, acyltransferase activity and mitochondrial activity.

**Figure 5.**
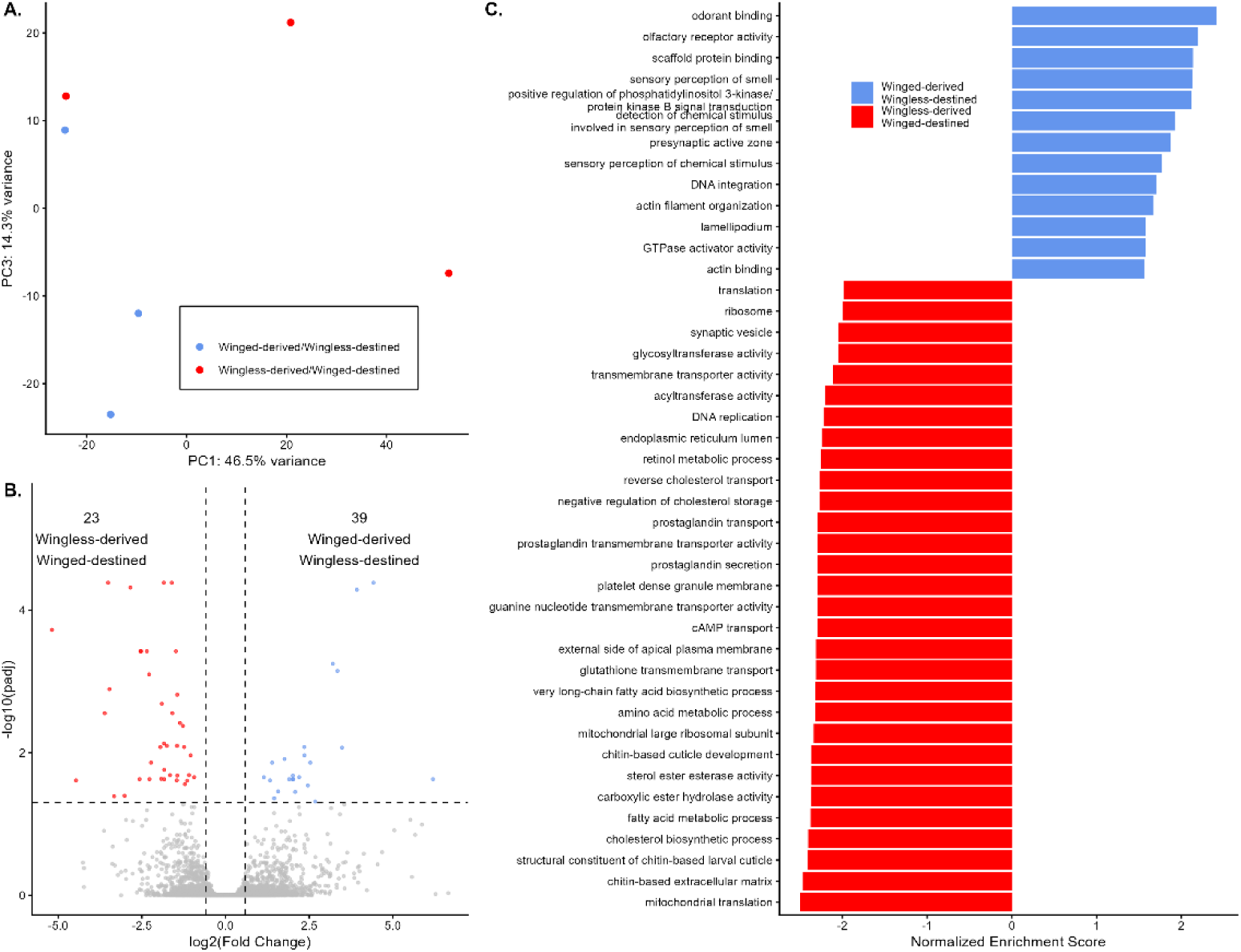
Gene expression differences associated with winged or wingless destined embryos. (A) Principal component analysis of 6 crowded embryo samples, using the 5,000 most variable genes. Points are colored by wing morph: wingless-derived/winged-destined samples (red) and winged-derived/wingless-destined samples (blue). (B) Volcano plot of differentially expressed genes between the two embryo types. Points are colored by wing morph: wingless-derived/winged-destined biased genes (red) and winged-derived/wingless-destined biased genes (blue). (C) Significantly enriched pathways identified by GSEA (adjusted p-value < 0.05). Data colored by wing morph: all wingless-derived/winged-destined embryo biased pathways are shown in red, and top30 winged-derived/wingless-destined embryo biased pathways are shown in blue.

### Promoter accessibility associates with gene expression

Using paired ATAC-seq and RNA-seq data, we examined whether genes with promoter-like OCRs exhibit higher expression than those without them. We found that genes with promoter accessibility exhibited significantly higher expression (Figure 6A), consistent with active transcriptional regulation. We also found a significant positive correlation between chromatin accessibility at the promoter region and gene expression (mean observed R = 0.065 vs. randomized null hypothesis R = 9.2 x 10^-4^, p = 0.026, Figure 6B). Although modest in magnitude, this positive shift indicates that accessibility mildly increases expression genome-wide, supporting the regulatory relevance of the ATAC-seq data.

**Figure 6.**
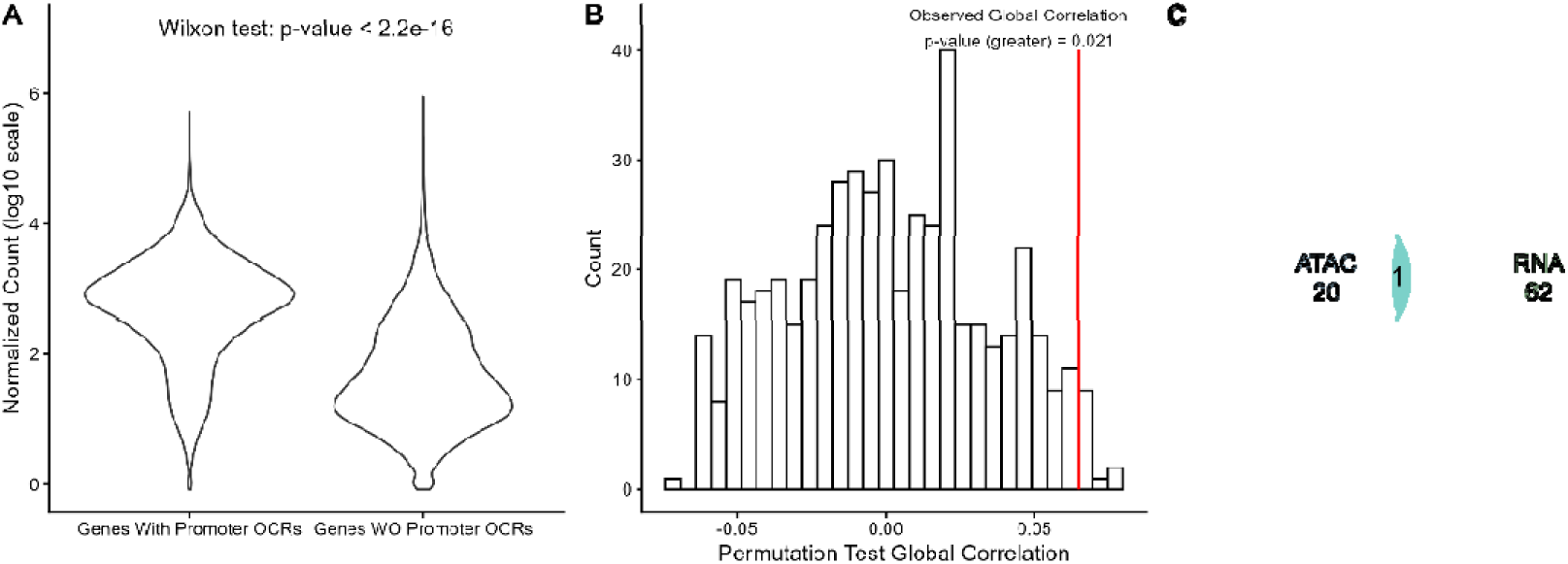
Integration of chromatin accessibility and gene expression. (A) Comparison of gene expression levels between genes with promoter-associated open chromatin regions (promoter-like OCRs) and genes without promoter-like OCRs. Statistical significance was assessed using the Wilcoxon rank-sum test. (B) Histogram of the global permutation correlation between promoter-region accessibility and gene expression. Observed correlation indicated in the red line. (C) Overlap between genes near DARs and differentially expressed genes (DEGs).

For 17 DARs, we retrieved 20 genes. Compared with the DEGs, we observed only 1 overlap (Figure 6 C). This gene is uncharacterized and its homolog in Drosophila melanogaster is annotated as 1-phosphatidylinositol 4,5-bisphosphate phosphodiesterase epsilon-1.

## Discussion

Here we present the first genome-wide chromatin accessibility map generated by ATAC-seq in pea aphids, characterizing OCRs across adult brains and late-stage embryos. We find that OCRs are broadly distributed across promoter and distal genomic regions, overlap substantially with computationally predicted enhancers, and are enriched for binding motifs of relevant transcription factor families. Chromatin accessibility is positively correlated with gene expression genome-wide, and tissue identity is the overwhelming driver of accessibility variation. Together, these results establish a regulatory genomics resource for pea aphid and provide a framework for dissecting *cis*-regulatory control of gene expression in this species.

The clearest finding of this study is the extent to which tissue identity dominates the chromatin accessibility landscape. Tissue alone accounted for 85% of the variance along the first principal component, and direct comparison between brain and embryo identified 19,511 DARs (52.6% of the 37,125 total)—a magnitude that far exceeded the chromatin differences associated with any other experimental factor. This result is consistent with the general principle that tissue-specific gene expression programs are accompanied by tissue-specific chromatin accessibility (Cusanovich et al. 2018). Moreover, the identities of genes near DARs are biologically coherent: brain-biased DARs are enriched near genes annotated with neural functions (axon, synapse, neuron projection), whereas embryo-biased DARs are predominantly enriched near genes involved in metabolic processes.

Compared to the extensive tissue-level chromatin differences, differences associated with wing morph or crowding treatment were modest. We identified 976 DARs between winged- and wingless-destined embryos and four between adult winged and wingless brains; no treatment-associated DARs were detected in any comparison. Together, these observations suggest that morph-associated regulatory divergence is subtle at this stage and that relatively few changes in the regulatory landscape may be sufficient to commit cells to alternative morph fates. This finding is also unsurprising given that we selected this embryonic stage because it falls within the earliest known window of wing morph differentiation (Ogawa and Miura 2013; Grantham et al. 2020). We predict that profiling later developmental stages would reveal greater divergence, consistent with the expansion of morph-associated transcriptional differences as development proceeds (Saleh-Ziabari et al., 2025).

Despite the modest morph-associated chromatin signal, the paired ATAC-seq and RNA-seq analysis revealed a significant positive genome-wide correlation between chromatin accessibility and gene expression. Genes with promoter-associated OCRs were also expressed at significantly higher levels than those without. These results validate the regulatory relevance of the identified OCRs and are consistent with the established mechanistic link between promoter accessibility and transcriptional output (Klemm et al. 2019). The limited overlap between DAR-proximal genes and DEGs is not surprising given that DARs are mostly found in intergenic regions and may regulate genes beyond a 30 kb proximity window, and that the modest magnitude of both chromatin and expression differences as well as low ATAC-seq library depth after accounting for endosymbiont contamination reduces power to detect concordant changes in both data types simultaneously. Together, these analyses demonstrate that the OCRs identified here capture functionally relevant regulatory regions.

A notable practical challenge in generating these data was the high abundance of reads derived from aphid endosymbionts, particularly *Spiroplasma* in brain samples and *Buchnera* in embryo samples. This likely reflects the preferential insertion of Tn5 into bacterially-derived DNA, which lacks nucleosome-mediated chromatin compaction. Similar enrichment of organellar or microbial DNA in ATAC-seq studies has previously been noted (Cantin et al. 2024). Importantly, once symbiont-derived reads were computationally removed, the remaining aphid-mapping libraries showed hallmarks of high-quality ATAC-seq data, including strong nucleosome-free fragment enrichment at transcription start sites. The challenges we encountered in adapting ATAC-seq to this system are consistent with the broader difficulties of implementing the method in non-model arthropods, as summarized in (Erdoğan et al. 2025). Future work could further improve resolution by increasing sequencing depth and more effectively removing endosymbiont contamination and thereby improving the sensitivity for detecting regulatory elements and subtle chromatin accessibility differences.

The consensus set of 37,127 OCRs identified here is broadly consistent in scale with ATAC-seq datasets from other insects, including *Drosophila melanogaster* (Blythe and Wieschaus 2016) and *Apis mellifera* (Lowe et al. 2022). Together with the whole-body FAIRE-seq chromatin accessibility data from (Richard et al. 2017), which linked X chromosome accessibility to dosage compensation in male and female pea aphids, the OCR atlas presented here substantially expands the regulatory genomics toolkit available for the pea aphid. As genomic and functional resources continue to expand in pea aphids and related species, these chromatin accessibility data will provide a foundation for mechanistic studies of the gene regulatory programs underlying the many remarkable biological phenomena in this group.

## Acknowledgements

Research reported here was supported by National Institute of General Medical Sciences of the National Institutes of Health under award number R35GM144001. We thank the University of Rochester Genomics Research Center for generating the sequencing data. We also thank Karl M. Glastad for discussions and suggestions regarding ATAC-seq sample preparation and data analysis.

---

**Figure S1.**
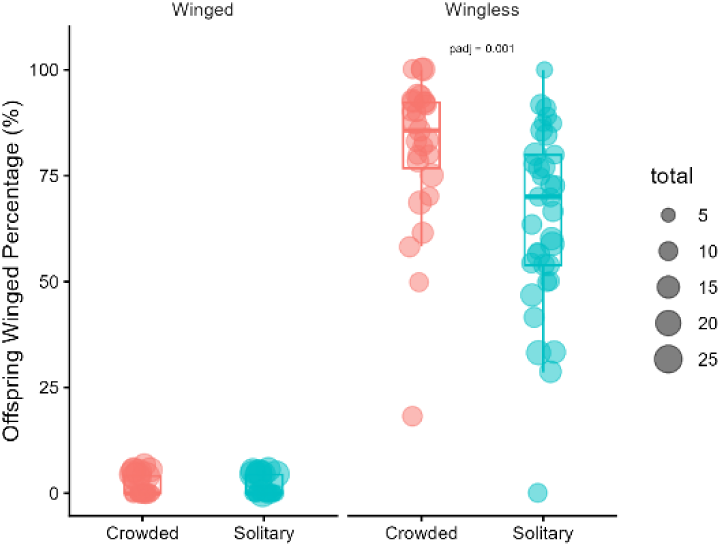
Wing morph phenotype across treatment and adult types. Winged adults (left panel) produce predominantly wingless offspring. For wingless adults (right panel), crowding induced a modest but significant increase in winged offspring compared to solitary conditions (Wilcoxon test, adjusted p-value = 0.001). The size of points represents the total number of offspring assessed for each replicate.

**Figure S2.**
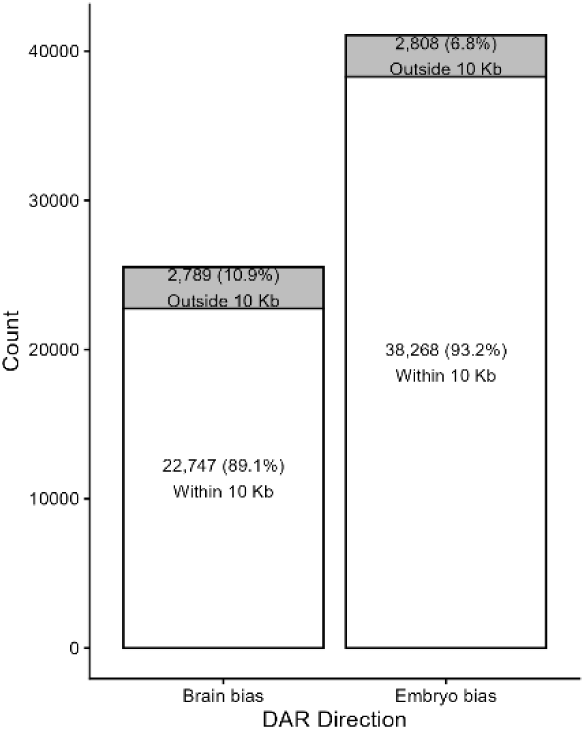
Number of genes with close proximity to DARs.

**Figure S3.**
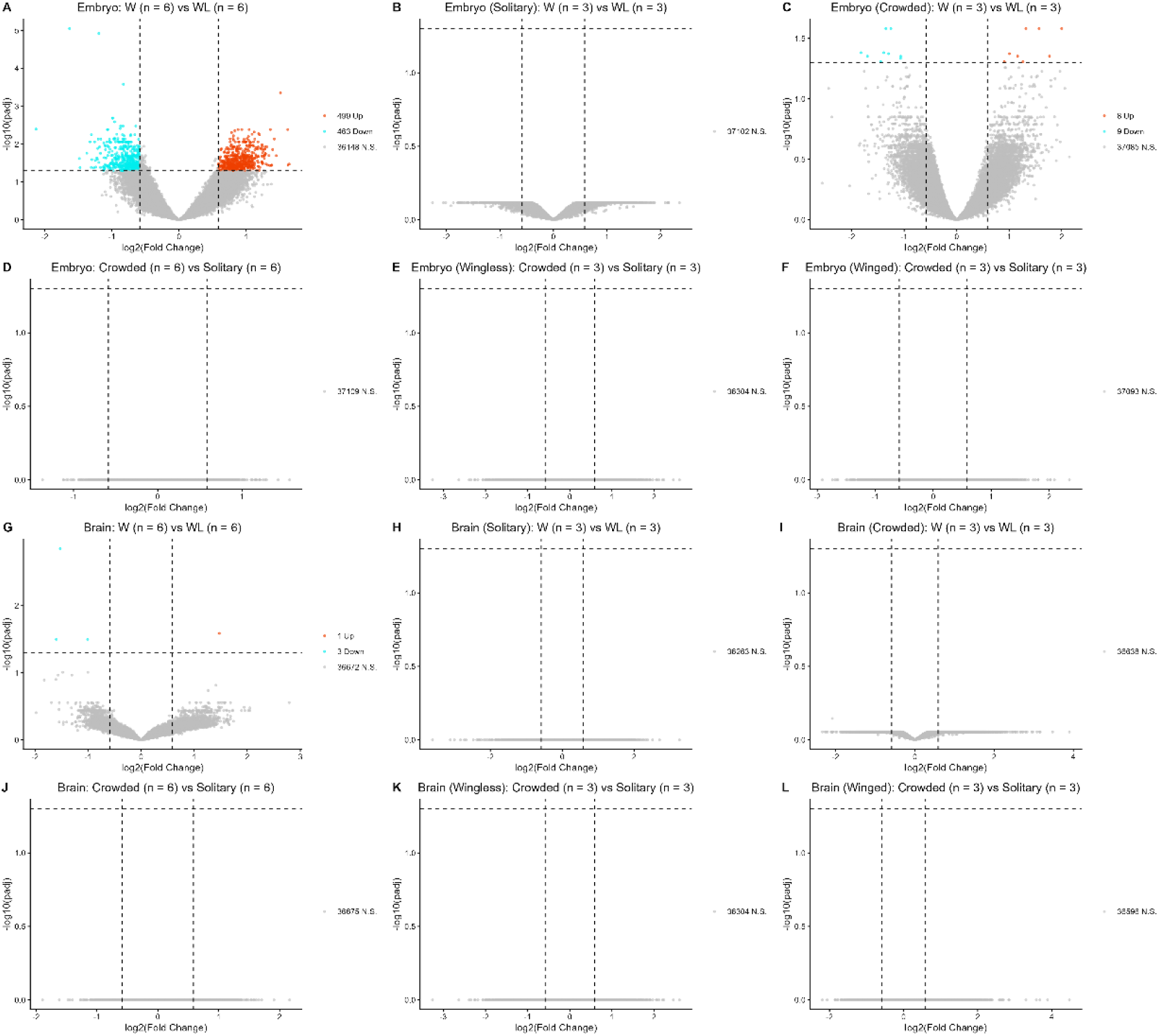
Volcano plot showing DARs for all pairwise comparisons.

**Table S1. Detailed supplementary information.**

