## Supplementary figures and images for "A chromatin accessibility map of pea aphid brain and embryo identifies tissue-specific regulatory elements"

### Figure S1

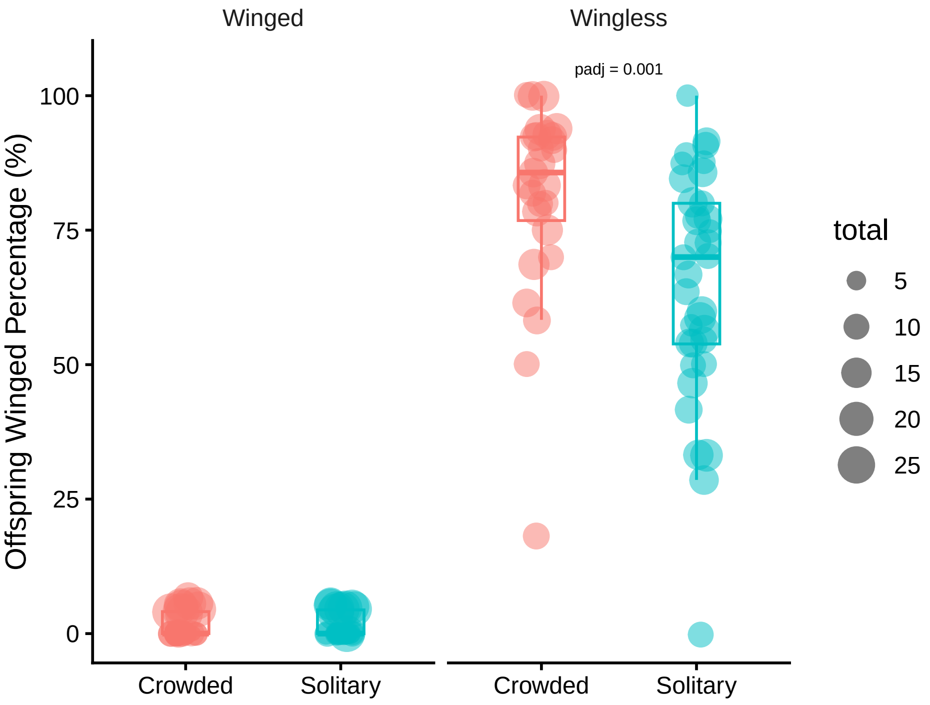

### Figure S2

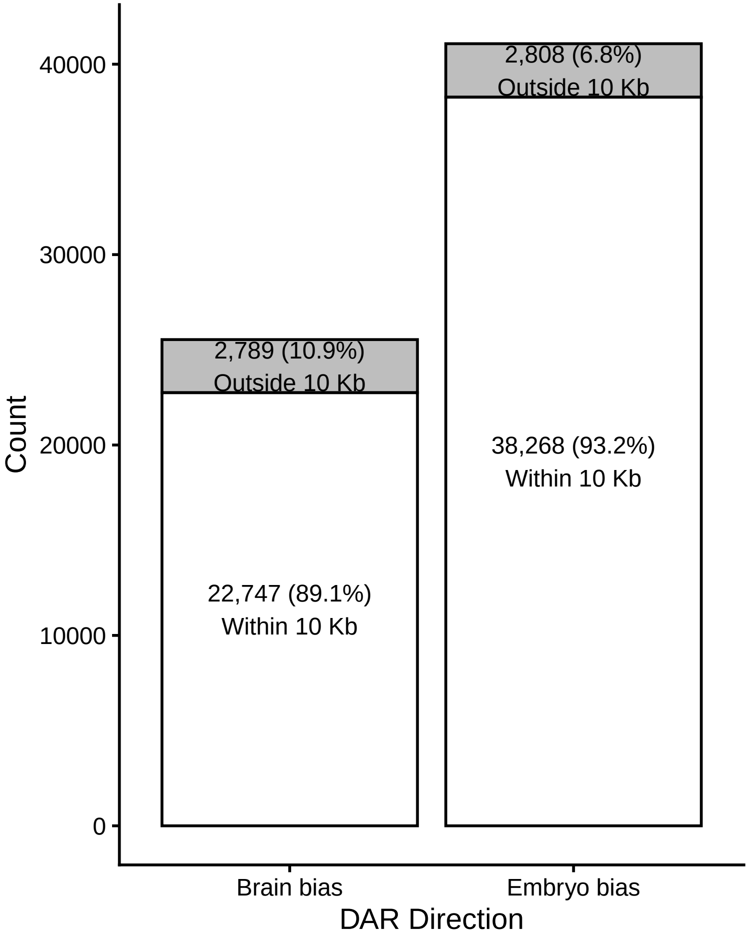

### Figure S3

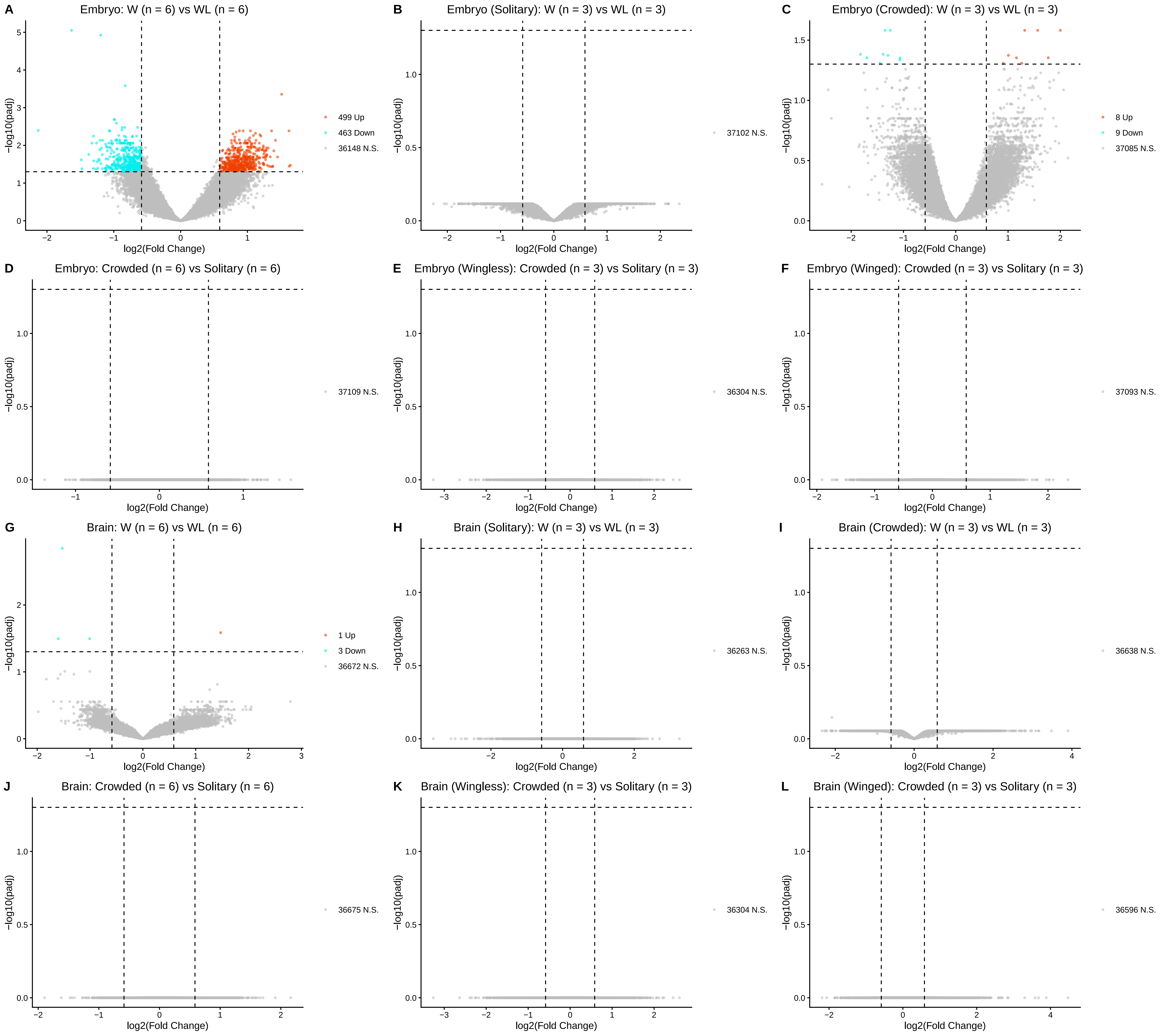
